# An improved graph representation learning method for drug-target interaction prediction over heterogeneous biological information graph

**DOI:** 10.1101/2022.06.30.498357

**Authors:** Bo-Wei Zhao, Xiao-Rui Su, Zhu-Hong You, Peng-Wei Hu, Lun Hu

## Abstract

The prediction task of the relationships between drugs and targets plays a significant role in the process of new drug discovery. Computational-based strategies for predicting drug-target interactions (DTIs) are regarded as a high-efficiency way. Moreover, recent studies adopted a graph neural network (GNN) to discover underlying DTIs and achieved better performance. Although these inductive methods can straightway learn biomolecules’ latent representations, they have an over-smoothing phenomenon in the course of obtaining the rich neighborhood information of each node in the biological information network, which further leads to a consistent feature representation of each node. To address the above issues, a novel model, called iGRLDTI, is proposed to precisely identify new DTIs based on an improved graph representation learning strategy. Specifically, iGRLDTI first constructs a biological information graph (BIG) by calculating the biological knowledge of drugs and targets with the relationships between them. Then, an improved graph representation learning strategy is designed to capture the enriched feature representations of drugs and targets. Finally, the Gradient Boosting Decision Tree classifier is applied to predict potential DTIs. Experimental results demonstrate that iGRLDTI yields better performance by comparing it with other state-of-the-art models on the benchmark dataset. Besides, our case studies denote that iGRLDTI can successfully identify unknown DTIs according to the improved feature representations of drugs and targets.

## Introduction

Identifying the relationship between drug and target is a crucial stage in drug discovery and development, while traditional experiments are usually performed by *in vitro* experiments. Instead of that, *in silico* experiments can be used as an alternative to discover new drugs, which can reduce the consumption of money and improve the efficiency of drug production [1]. Generally speaking, *in silico* screening methods are classified into two categories, including docking simulations and machine learning methods [2].

The former is on the base of the drug molecule and the target three-dimensional structure to discover a binding site suitable, which step is not only time-consuming but also subject to target structure information [3]. Taking the G-protein-coupled receptors (GPCR) as an example, there are relatively few that can be crystallized (orphan GPCR), which makes docking simulation experiments difficult to implement [4].

Therefore, most of the studies consider drug-target interactions (DTIs) are in the light of chemogenomic, which can link compounds and target proteins to confirm new compounds and targets [5]. Also, chemogenomic methods are essentially based on the similarity criterion between compounds and targets, and thus machine learning methods are more suitable [6, 7].

In recent years, machine learning-based methods have been widely and successfully applied to DTI prediction [8], which mainly contains four steps: (i) data collection and preprocessing; (ii) feature extraction; (iii) classification and prediction; (iv) experiments validation. In practice, based on the differences in the feature extraction methods for drug compounds and target proteins, the DTI prediction methods can be categorized into three types: network-based approaches, matrix factorization-based approaches, and deep learning-based approaches [8, 9]. The network-based approach integrates the drug-target network and the drug compound and target protein-related network to obtain their characteristics for the prediction task of DTIs. For instance, Wang et. al [10] proposed a triple-layer heterogeneous network model for drug repositioning, which combined three types of nodes, i.e., diseases, drugs, and targets. Luo et al. [11] presented a novel model with a network integration pipeline, called DTINet, which integrated multiple information from the heterogeneous network. DTINet first integrated a diverse drug-related network to construct a heterogeneous network, and then a compact feature learning method is applied to learn the low-dimensional vector representation with the topological properties of each node. Wan et. al [12] developed a nonlinear end-to-end learning model, namely NeoDTI, by integrating multiple information from heterogeneous network data to learn the network representations of drugs and targets for DTI prediction. NeoDTI first integrates neighborhood information of nodes according to information passing and then a network topology-preserving learning procedure is utilized to extract the representations of drugs and targets. However, they ignore the attribute information of the biological data itself when obtaining the diffusion state of the node in the biological network, which may lead to inaccurate DTI predictions.

The matrix factorization-based approach is as a way based on an association matrix and similarity matrix. It assumes that there are certain relationships between drugs, targets, and diseases, and can effectively discover their relationships. Liu et. al [13] used a neighborhood regularized logistic matrix factorization method to represent drugs and targets in the shared low-dimensional latent space and produce the probability of drug-target interaction. Yang et. Al [14] designed a multi-similarities bilinear matrix factorization model to extract the latent features of drug compounds and diseases, and then infer unknown associations. Although useful, like network-based methods, it fails to use molecular biological knowledge and the effects of emerging drugs and targets are unsatisfactory.

The deep learning-based approach has received extensive attention in the field of drug research due to its outstanding performance in computer vision, natural language processing, and other fields [15-17]. Wang et. al [18] proposed a deep learning computational method for predicting DTIs, which mainly used the deep learning stacked autoencoder to train the drug molecular structure information and protein sequence information. DeepFusionDTA [19] as a deep neural network ensemble model with various information analysis modules is applied to detect DTIs. However, the above methods either ignore the biological attribute of drugs and targets or fail to learn the topology structure of DTIs. Recently, graph neural network-based methods are widely used in bioinformatics due to the better fusion of the biological attribute and topology structure [20-24]. For example, IMCHGAN [25] first constructed diverse meta-paths about drugs and targets, and then a two-level neural attention mechanism is introduced to learn the representations of drugs and targets. MultiDTI [26] combined various association information and the sequence information of drug and target into a common space to predict new DTIs. However, the above methods typically more or less fail to provide satisfactory prediction performance. For example, DTINet more focus on network structure while ignoring the biological data itself attribute, which is difficult to fully capture the representation of biomolecules to accurately predicting DTIs. Furthermore, recent methods have also started simultaneously taking into account biological attributes and network structures in the biological network, such EEG-DTI [27] and IMCHGAN [25] have achieved better prediction results. Yet, due to GNN itself exists the limit with over-smoothing. As a result, the learned feature representations of all nodes may tend to be the same rather than partially similar.

To overcome this issue, we consider optimizing neighborhoods’ information aggregation operation by calculating the level of neighbor nodes of each node. Specifically, we first calculate the impact of neighboring targets on each drug to build a degree matrix with neighbor node, and then by setting a variate *k*to adapt the number of neighbors’ information aggregation iterations. In doing so, we can obtain more high-quality feature representations to predict DTIs.

In this paper, we propose a novel model, termed iGRLDTI, to improve the accuracy of DTI prediction based on improved graph representation learning strategies. More specifically, iGRLDTI first calculates the biological knowledge of drugs and targets including drug molecular structure information and target sequence information, to obtain a BIG with the relationships between drugs and targets collected. After that, an improved graph representation learning strategy is constructed to capture the optimum feature representations of drugs and targets. In particular, the influence between drugs and targets is calculated to control the operation of nodes’ aggregate neighbor information based on the graph representation learning, thereby obtaining excellent feature representations. Finally, the Gradient Boosting Decision Tree (GDBT) classifier is introduced as a classifier tool to complete the DTI prediction task. Comprehensive experiments demonstrate that iGRLDTI yields significant performance on the benchmark dataset when it compares several state-of-the-art methods. The main contributions of this work are as follows:

(1) By calculating the influence between drugs and targets, a degree matrix is constructed to further capture the salient feature representations of drugs and targets.

(2) A novel BIG-based DTI prediction method, namely iGRLDTI, is proposed to precisely discover novel DTIs based on improved graph representation learning strategies.

(3) Experimental results demonstrate that iGRLDTI performs better performance than several state-of-the-art methods on the benchmark dataset. Furthermore, the case studies indicate that iGRLDTI yields more practicable performance for identifying new associations between drugs and targets.

## Methods

### Model structure

iGRLDTI mainly consists of three steps: (i) constructs a BIG by obtaining the biological features of drugs and targets with the biological knowledge collected and the drug-target associations. (ii) calculates the influence between drugs and targets to optimize the operation of nodes aggregate neighbor information in the course of learning feature representations for drugs and targets; (iii) discovers novel DTIs by the GDBT classifier. The schematic diagram of iGRLDTI is presented in Figure 1. In particular, we construct the mathematical formulations of the following steps.

**Figure 1.**
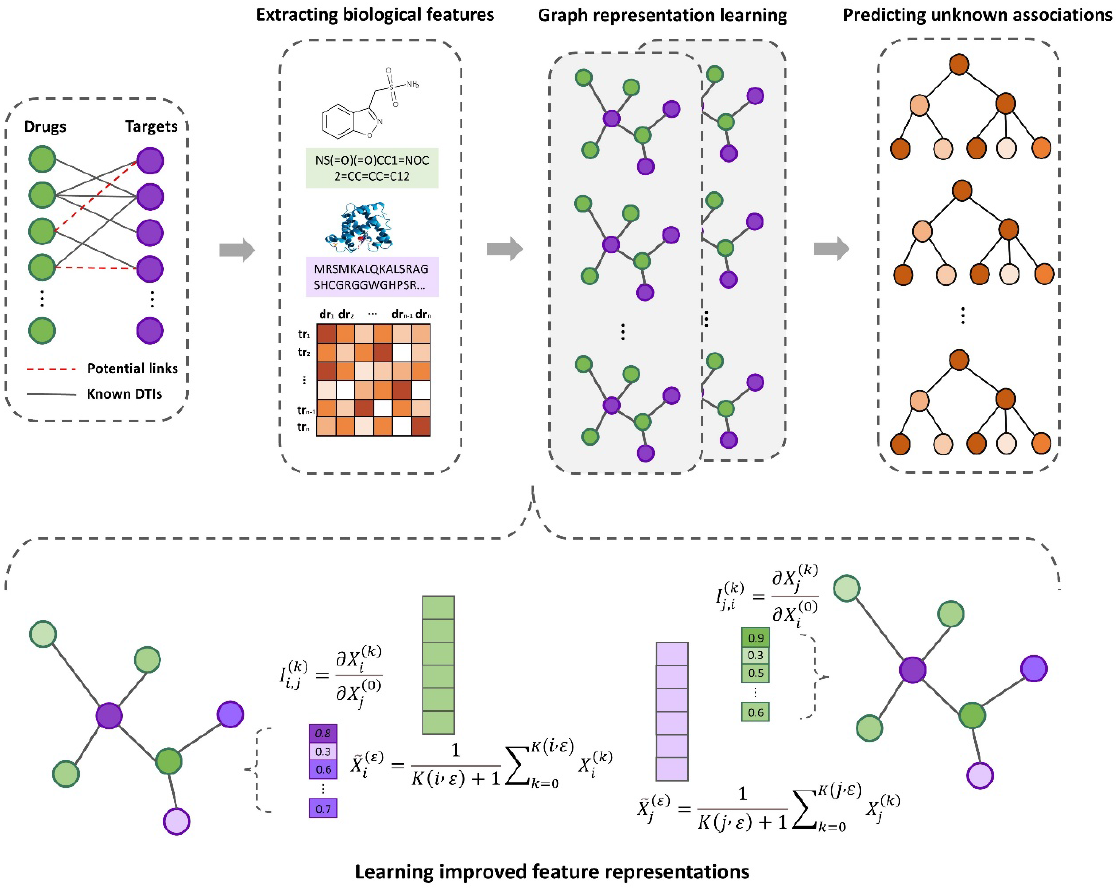
The schematic diagram of iGRLDTI.

Definition 1: Given a biological information graph (BIG), which contains three elements, i.e., BIG={*V,C,E*}, *V*={*V*^*dr*^,*V*^*tr*^}signifies all nodes (|*V*|) in the BIG and all drugs (*V*^*dr*^) and all targets (*V*^*tr*^}). *C* denotes the biological knowledge of drugs and targets. *E* represents the relationships between drugs and targets, |*E*| is the number of DTIs. Moreover, the number of drugs and targets is denoted as *n* and *m* respectively.

Definition 2: Regarding the drug molecular structure information and target sequence information are represented as *C*^*dr*^ ∈ *ℜ*^*n*×*d*^ and *C*^*tr*^ ∈ *ℜ*^*m*×*d*^ respectively. The biological feature matrix is denoted as 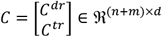. Besides, the adjacent matrix of the drug-target network is regarded as *A* ∈ *ℜ*^|*V*|×|*V*|^.

Definition 3: Given a matrix *I* ∈ *ℜ*^|*V*|×|*V*|^ is expressed to collect the influence between drugs and targets, and the variate *k* is set as the number of iterations in the graph representation learning.

### Extracting drug biological features

As regards the biological knowledge of drugs, we take advantage of the Simplified Molecular Input Line Entry System (SMILES) [28], which is a linear symbol for the representation of molecular reactions, to represent drug molecule information (*C*^*dr*^). Thus, we collect the SMILES of each drug from the DrugBank database [29], and then calculate the SMILES of each drug by the RDKit tool [30] to obtain the drug’s biological feature *c*^*dr*^.

Since *C*^*dr*^ is high-dimensional, we attempt to construct an auto-encoder to reduce its dimensionto *d*-dimensional (i.e *d* = 64). At last, all drug features are defined as 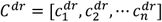.

### Extracting protein biological features

At the present stage of computational methods, the biological knowledge of proteins is mainly four classes, i.e., genomic information, protein structures, protein sequences, and GO [31-33]. In which, the protein sequence is widely applied to protein-related prediction tasks [34-36]. In this work, we collect the sequence information (*C*^*tr*^) of each protein as the biological knowledge of them, and then k-mer algorithm is used to extract all protein features 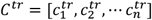 by calculating the frequency of protein sequence substrings. Note that the k-mer algorithm is set as 3-mer, which is a 64-dimensional vector (i.e *d* = 64) to represent the features *c*^*tr*^ of each target.

### Learning improved feature representations

Exiting research generally treats network topology and biological knowledge as nodes’ feature representations[22, 37]. For instance, IMCHGAN [25] and GVDTI [38] use different GNN such GAT [39] and GCN [40] to treat the biological information of drugs and targets. However, these algorithms need outstanding prior biological knowledge to learn hopeful representation by GNN, because there is a certain degree of over-smoothing phenomenon, that is, the obtained representations of the nodes may tend to be consistent, which makes it difficult to distinguish the downstream classification tasks. Generally, the representations of drugs and targets by means of the following aggregation:

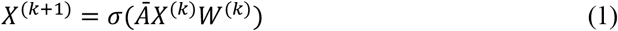

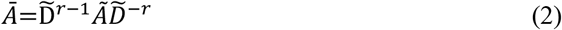

where *Ã* is the adjacency matrix *A* with self-loops, and 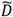 is a degree matrix of the adjacency matrix *Ã, r* = [0,1]. *k* is the layer number of the neural network. *W* is a weight matrix, and *σ* is an activation function. To make it easy to get the smoothed feature after *k*-iterations, propagation [41, 42]. Let us assume *σ*(·)and *W* are an identity function and an identity matrix respectively, and then Equation (1) can simplify as following form.

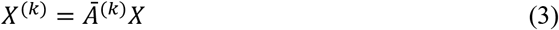

Moreover, after the limitless propagation, the continually smoothed features of the node can be represented as:

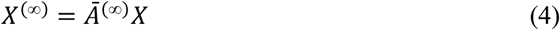

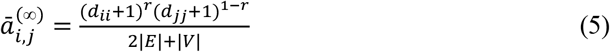

where 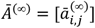 represent the latest adjacency matrix with smoothed, *d*_*ii*_ and *d*_*jj*_ are the degrees of the drug 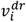 and the target 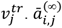 represent the influence of the drug 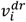 and the target 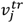 Hence, the feature representations of drugs and targets when *k*-iterations later are defined as:

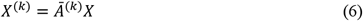

Inspired by [43], we measure the influence of the drug 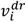 on 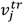 by calculating how much a change in the biological feature 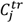 of 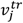 affects the feature representation of 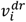 after *k*-iterations. Given a degree matrix *I* to present the influence between drugs and targets.

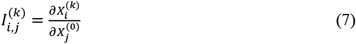

where 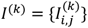, According to Equation (4), Equation (6), and Equation (7), we can define *I*^(*k*)^ and *Ĩ*^(∞)^ as:

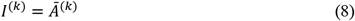

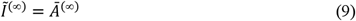

To realize the comparison between the feature representation of 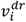 $ after *k*–iterations propagation and after infinite propagation, a parameter *ε* (*ε* > 0) is introduced to confirm the minimal value of *k*-iterations needed for the influence of neighbor 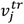 on 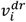 when the feature representation *X* converges to the stationary distribution. Given a function *K* to collect the above relationship by:

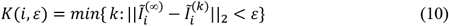

where || · ||_2_ is two-norm and *K*(*i,ε*) >. After that, the biological feature *X*_*i*_ of 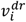 can be converted as follow:

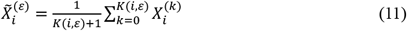

Moreover, a fully connected layer of neural networks is applied to obtain the final feature representations *X*^*dr*^ ∈ *ℜ*^*n*×*d*^ of drugs by:

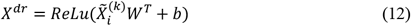

In the above equation, *ReLu*(·) is the activation function, *W* ∈ *ℜ*^*d*×*d*^ is the weight matrix from the input layer to the hidden layer and *b* is the bias. Similarly, we can also obtain *X*^*tr*^ as the final feature representations of targets. Hence, a (*n* + *m*) × *d* matrix *X* = [*X*^*dr*^,*X*^*tr*^]^*T*^ is used to denote the representation vectors of drugs and targets in a more concise manner.

### Predicting novel DTIs via GBDT

To achieve the prediction of drugs and targets, iGRLDTI selects a typical machine learning classifier with ensemble learning, namely the Gradient Boosting Decision Tree (GBDT) classifier, which trains features on multiple Decision trees and takes the conclusions of these trees as the final prediction results. Since the association prediction belongs to a binary classification problem, we construct the classification features required as *H* = [*X*^*dr*^,*X*^*tr*^] ∈ *E*. A matrix *R* is used to collect the prediction results of iGRLDTI, in which *R* only contains 0 and 1 to represent disconnect or connect. The complete procedure of iGRLDTI is described in **Algorithm 1**.

## Results

### Evaluation criteria

To better demonstrate the performance of iGRLDTI, we select the benchmark dataset to rely on Luo’s research [11], including 1923 drug-target interactions, 549 drugs, and 424 targets. After that, AUC and AUPRC are used as standardized evaluation indicators, in which the AUC is an area under the receiver operating characteristic (ROC) curve and AUPRC is an area under the precision-recall (PR) curve. Besides, several indicators are introduced as a way to evaluate the prediction performance of iGRLDTI, including Precision, Recall, and F1-score, and they are calculated as follows:

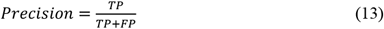

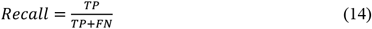

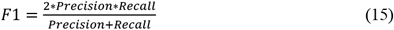

where *TP* and *TT* are the numbers of predicted true positive and negative samples, *FP* and *FT* are predicted false positive and negative samples. In the experiment, ten-fold cross-validation (CV) is used as a scheme for the performance evaluation of iGRLDTI. More specifically, we randomly divide the benchmark dataset into 10 of the same number of subsets, each fold selects 9 subsets of them as the training sets and the remaining subset as the test set. Moreover, the number of positive and negative samples is consistent for each training and test.

#### Algorithm 1

**The complete procedure of iGRLDTI**

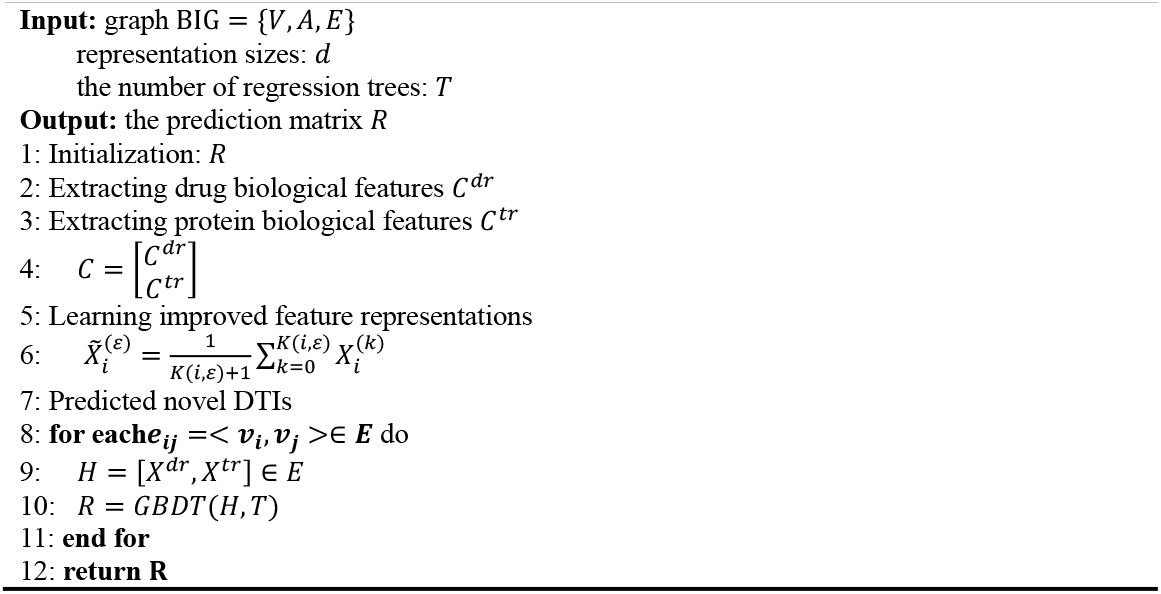

### Comparison with state-of-the-art models

To accuracy evaluation the performance of iGRLDTI, we compare it with four state-of-the-art methods proposed for DTI prediction, i.e., DTINet [11], NeoDTI [12], IMCHGAN [25], and MultiDTI [26]. The predicted experiment results of these methods under 10-fold CV are displayed in Table 1. We note should that iGRLDTI achieves the highest performance for DTIs in light of the AUC, AUPR, and F1 values of all methods. Among which, the AUC, AUPR, and F1-score of iGRLDTI higher than the second-highest methods are 0.7%, 2.3%, and 3%, respectively. In this regard, iGRLDTI can improve the feature representations of drugs and targets by optimizing neighbor information aggregation for improved accuracy in the course of DTI prediction.

**Table 1.** Comparison with state-of-the-art models on benchmark dataset.

| Methods | AUC | AUPR | F1-score |  |  |
| --- | --- | --- | --- | --- | --- |
|  |  |  | Pre | Rec | F1 |
| DTINet | 0.914 | 0.932 | **1.0** | 0.048 | 0.090 |
| NeoDTI | 0.957 | 0.874 | 0.928 | 0.716 | 0.808 |
| IMCHGAN | 0.957 | 0.904 | NA | NA | NA |
| MultiDTI | 0.961 | 0.947 | 0.876 | 0.882 | 0.868 |
| iGRLDTI | **0.968** | **0.970** | 0.900 | **0.896** | **0.898** |

Except for the analysis of these values, we also note that the values of some indicators appear to be satisfactory, and further in-depth analyses of the reason for that phenomenon are unbalanced positive and negative samples. Take NeoDTI as an example, which uses 10 times as many positive samples as unknown relations of negative samples during the training and test. In doing so, not merely its prediction results are biased to the classification with more samples, that is, the number of negative samples is more predicted, but also the evaluation indexes of the model could be won higher, such as AUC. However, a model usually needs to be able to discriminate which samples are true positive samples and which are false negative samples in a biomolecule associations prediction task. Regarding the perfect precision score and the lowest recall score of DTINet, the reason is due to its predicted results containing too many false-negative samples and scarce true positive samples, thereby reducing the F1-score. In summary, iGRLDTI can better absorb neighbor features to improve its feature representation and further enhance its predictive ability.

### Comparison of different feature extract methods

Regarding the influence of different features for DTI prediction, we also perform other two variants on the base of iGRLDTI, i.e., iGRLDTI-A and iGRLDTI-G. Specifically, iGRLDTI-A merely considers drug molecule structure information and protein sequence information, and iGRLDTI-G learns the feature representations of drugs and targets based on a general graph convolution network. On the whole, these two variants also use the GDBT classifier with the same hyperparameters to predict novel DTIs under 10-fold CV. In Table 2, we have compared the evaluation indexes of iGRLDTI-A, iGRLDTI-G, and iGRLDTI.

**Table 2.**
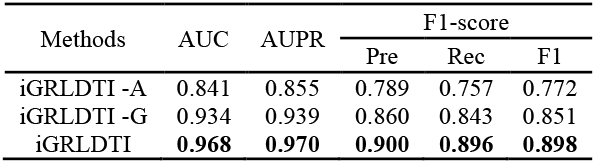
Performance of different feature extract methods.

Additionally, it is noteworthy that iGRLDTI-A achieves the worst score compared with another two models. In doing so, only the biological knowledge is difficult to build a model with a strong performance for DTI prediction. Since *C* is based on features collected manually (i.e., manual download and manual calculation), which involves a phenomenon such as missing biological knowledge of some nodes and errors caused by calculations and further leads to the fact that the features cannot be accurately distinguished in the classifier. Take the drug as an example, only 536 of these drugs obtained SMILES and the remaining could not find their SMILES from the database. Meanwhile, the molecular structure of the drug 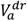 and 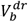 is more similar, but there may have very different SMILES strings [44]. Thus, only relying on biological knowledge may not be accurately distinguished unknown relationships between drugs and targets. And that’s why iGRLDTI-G could be reached quite high change. In particular, iGRLDTI-G performs better by 9.3%, 8.4%, 7.1%, 8.6%, and 7.9% than iGRLDTI-A in terms of AUC, AUPR, Pre, Rec, F1-score, respectively. Indeed, the biological knowledge of drugs and targets can be enhanced by the graph neural network, and then the powerful combination of network information and neighbor information can better improve the expressive ability of feature representations of drugs and targets. iGRLDTI achieves better performance when it learns the feature representations of drugs and targets than iGRLDTI-G, and further can effectively reduce over-smoothing in the course of aggregating neighbor nodes information.

### Predicting novel DTIs

To demonstrate the practical ability of iGRLDTI for identifying unknown drug-protein interactions, we conduct case studies on the benchmark dataset. This experiment yields a high achievement through training all known samples. In particular, we use all known DTIs as training samples to obtain iGRLDTI, and then iGRLDTI is applied to discover all unidentified DTIs. We collect and verify the top 20 candidate samples and the detailed results as presented in Table 3. One should note that all candidates are proved by the DrugBank dataset. This result indicated that iGRLDTI has an outstanding ability to predict potential drug-related targets and can be used as a promising tool to gain new insight into the task of novel DTI prediction by obtaining their smoothed feature representations.

**Table 3.** Top 20 predicted results by iGRLDTI.

| Drug ID | Drug Name | Protein ID | Protein Name | Evidence |
| --- | --- | --- | --- | --- |
| DB00909 | Zonisamide | O43570 | Carbonic anhydrase 12 | DrugBank |
| DB01110 | Miconazole | Q14500 | ATP-sensitive inward rectifier potassium channel 12 | DrugBank |
| DB00909 | Zonisamide | Q99250 | Sodium channel protein type 2 subunit alpha | DrugBank |
| DB00594 | Amiloride | P19634 | Sodium/hydrogen exchanger 1 | DrugBank |
| DB01159 | Halothane | P48051 | G protein-activated inward rectifier potassium channel 2 | DrugBank |
| DB00661 | Verapamil | O95180 | Voltage-dependent T-type calcium channel subunit alpha-1H | DrugBank |
| DB00594 | Amiloride | P19801 | Amiloride-sensitive amine oxidase [copper-containing] | DrugBank |
| DB01159 | Halothane | O60391 | Glutamate receptor ionotropic, NMDA 3B | DrugBank |
| DB01159 | Halothane | P48549 | G protein-activated inward rectifier potassium channel 1 | DrugBank |
| DB00909 | Zonisamide | O43497 | Voltage-dependent T-type calcium channel subunit alpha-1G | DrugBank |
| DB01268 | Sunitinib | P17948 | Vascular endothelial growth factor receptor 1 | DrugBank |
| DB00909 | Zonisamide | P00918 | Carbonic anhydrase 2 | DrugBank |
| DB00398 | Sorafenib | P17948 | Vascular endothelial growth factor receptor 1 | DrugBank |
| DB01224 | Quetiapine | P28335 | 5-hydroxytryptamine receptor 2C | DrugBank |
| DB01268 | Sunitinib | P09619 | Platelet-derived growth factor receptor beta | DrugBank |
| DB00398 | Sorafenib | P15056 | Serine/threonine-protein kinase B-raf | DrugBank |
| DB01268 | Sunitinib | P36888 | Receptor-type tyrosine-protein kinase FLT3 | DrugBank |
| DB00398 | Sorafenib | P04049 | RAF proto-oncogene serine/threonine-protein kinase | DrugBank |
| DB01110 | Miconazole | Q13936 | Voltage-dependent L-type calcium channel subunit alpha-1C | DrugBank |
| DB00909 | Zonisamide | P21397 | Amine oxidase [flavin-containing] A | DrugBank |

## Discussion and conclusion

In this work, a novel prediction model, called iGRLDTI, is proposed to identify unknown DTIs. First, a BIG is constructed by the biological features of drugs and targets extracted from biological knowledge and the relationships between drugs and targets. Then, a modified graph representation learning method is developed by calculating the influence between drugs and targets to control the number of iterations to obtain the enhanced feature representations of drugs and targets. Finally, iGRLDTI combines the GDBT classifier to achieve the DTI prediction task. Experimental results demonstrate that iGRLDTI yields a high performance under 10-fold CV than other advanced DTI prediction methods, and the case studies show that it has a strong practical discovery ability for potential DTIs.

Although the experiment results have demonstrated the promising performance of iGRLDTI, there are still some limitations to be addressed in future work. On the one hand, we are concerned with the application of iGRLDTI on other biological molecules, such as protein-protein interactions prediction [32] and drug-disease interactions prediction [24]. On the other hand, we would like to improve the interpretability of iGRLDTI by addressing the constraint of graph representation learning [45].

## Notes

### Competing Interest Statement

The authors have declared no competing interest.

